# Field-tested *HaHB11* and *HaHB4* soybean exhibit increased grain number and heat tolerance at the reproductive stage

**DOI:** 10.1101/2024.09.26.615293

**Authors:** Jesica Raineri, Enrique Montero Bulacio, Mabel Campi, Margarita Portapila, María Elena Otegui, Raquel Lía Chan

## Abstract

Soybean is one of the primary sources of vegetable oil and protein worldwide. However, its yield improvement has lagged behind the other major crops. This study explored the potential of the sunflower transcription factor HaHB11 to enhance soybean yield and heat stress tolerance. We generated transgenic soybean plants expressing HaHB11 and evaluated their performance across four field trials. The HaHB11 plants showed a significant increase in grain number per plant compared to controls, which can be related to an increased number of nodes and pods per plant. Flowering dynamics analysis revealed delayed blooming and an increased number of flowers per node, leading to a higher pod set, particularly between nodes four and six. Principal component analysis across field trials identified temperature as a crucial factor influencing grain number, enhancing the differences exhibited by HaHB11 plants. The pollen from transgenic plants germinated better, and tubes were longer than controls under heat stress. Carbohydrate distribution analyses indicated differential allocation of nutrients, supporting the increased pod and grain set in HaHB11 plants. Additionally, vegetation indices can distinguish HaHB11 plants from controls in several developmental stages. These results indicated that HaHB11 enhances soybean yield under heat stress, becoming a promising technology for soybean improvement.

**Highlight:** Soybean transformed with the sunflower gene *HaHB11* was tested in the field for four campaigns, showing differential allocation of nutrients, increased number of nodes, pods, grains, and heat tolerance.

## Introduction

Soybean (Glycine max L. Merr.) is the major source of the world’s vegetable oil and a significant protein source (Graham and Vance, 2003). This crop was not part of the Green Revolution, and its yield increase, constrained by abiotic stress factors, has been slower than that of rice, maize, or wheat (Liu *et al*., 2020). Global soybean yield of the three main world producers increased linearly since 1960 at a rate of 1.50 % y^-1^ for Argentina, 1.88 % y^-1^ for Brazil, and 1.27 % y^-1^ for the United States (FAO, 2024). These trends were mainly due to optimized agricultural practices, new genotypes obtained by traditional breeding methods, molecular breeding assisted by molecular markers, mutation breeding, and biotechnology and transgenic approaches techniques (de Felipe *et al*., 2016; Anderson *et al*., 2019; Vogel *et al*., 2021). In the United States, selected crop varieties of maturity group III (MG III) experienced over time improved canopy light capture due to delayed senescence, reduced plant lodging, more efficient conversion of intercepted light into biomass, and increased partitioning of biomass to seeds (Koester *et al*., 2014). Similar results were obtained for seed yield, biomass production and partitioning for MGs III-V in Argentina (de Felipe et al, 2020) and MGs IV-VIII in Brazil (Todeschini *et al*., 2019). Currently, biotech soybean covers more than 90% of the soybean-cropped land in the USA and Argentina (http://www. isaaa.org/). This commercialized genetically modified (GM) soybean shows resistance to herbicide glyphosate or herbivore attack. Interestingly, in Argentina and Brazil, the global yield gain of this species decreased markedly since the release of Roundup Ready (RR) soybean in 1996 (from more than 2.1 % y^-1^ for the 1960-1995 period to less than 1.4% y^-1^ thereafter; FAO, 2024). This trend has been attributed to the large expansion of soybean-cropped land promoted by the RR technology, in Argentina particularly towards regions that are frequently exposed to water deficit and heat-shock events (Grau *et al*., 2005; Song *et al*., 2021). While many studies conducted in controlled environments, such as growth chambers or greenhouses, describe the performance of genetically modified soybean under abiotic stress (Li *et al*., 2017; Wang *et al*., 2017; Chen *et al*., 2019), scientific literature on abiotic stress tolerance and related traits under field conditions is scarce. It is well known that there is a long distance between a culture chamber experiment and a field trial, in which several factors act together. An additional problem is when plants were tested only in stressing conditions without evaluating their behavior in normal growth conditions. For example, for water deficit stress, most genes, used as transgenes in varied plant species, involved stomatal closure, which triggers survival but usually reduces biomass and seed yield under mild stress (Skirycz *et al*., 2011).

Yield has two components: the number of seeds and their individual weight, being the former the most relevant for soybean seed yield determination. Varieties released over the years showed a clear trend of setting more pods/seeds, with no clear trend in seed weight (de Felipe *et al*., 2016; Todeschini *et al*., 2019; Vogel *et al*., 2021). The number of seeds per plant is determined by the number of seeds per pod, the number of pods per node, and the number of fertile nodes per plant (Egli, 2023; Vogel *et al*., 2021). The pod setting closely depends on the number of flowers per node, flowering dynamics, and emergence synchrony (Bruening and Egli, 2000; Egli and Bruening, 2006). Synchronous flowering in a short period would reduce competition among pods during filling, leading to a lower rate of developmental arrest (Egli and Bruening, 2006). Day length and planting date strongly influence flowering dynamics. However, increased planting density, CO_2_ availability, and different fertilization treatments do not affect this parameter (Egli and Bruening, 2006).

Heat is one of the main types of abiotic stress affecting soybean crops. Temperatures above 35°C cause heat stress, leading to significant yield losses by affecting photosynthetic efficiency, flowering, and pod setting (Salem *et al*., 2007; Prasad *et al*., 2008; Nahar *et al*., 2016; Onat *et al*., 2017; Thomey *et al*., 2019). High temperatures during flowering reduce pollen production, germination, and pollen tube length (Koti *et al*., 2005; Djanaguiraman *et al*., 2013). When the stress occurs between R5 and R7, the number of set pods decreases, and the effective grain-filling period shortens, possibly due to accelerated senescence (Egli and Wardlaw, 1980; Djanaguiraman and Prasad, 2010). Hence, heat tolerance is a highly desirable trait for improving seed yield. Traditional breeding assisted with molecular markers and other tools contributed to achieving this goal. However, we are far away from the ideal soybean phenotype. Genetic modification could aid in developing heat stress-tolerant varieties, with transcription factors (TFs) being promising candidates for enhancing resilience (Mittler and Blumwald, 2010; Baillo *et al*., 2019; Jianing *et al*., 2022). TFs are proteins that regulate the expression of many downstream genes in molecular signaling pathways, and their presence, absence, or differential regulation deeply affects plants’ fitness. In plants, there are around 1500 different TFs, grouped into more than 80 families (Riechmann, 2002; González, 2016). Families were defined mostly according to the DNA binding domain of their members. Some families are numerous and were resolved in different clades of phylogenetic trees (González, 2016). Several are unique to plants and others are present in more kingdoms. Others exhibit divergent members that do not show high sequence identity with proteins from other organisms; those are thought to be related to neofunctionalization and stress mitigation mechanisms specific to plants. One of the specific plant families is the homeodomain-leucine zipper (HD-Zip), in which members combine a homeodomain (HD) with a leucine zipper (Zip). Both domains are present in TFs from other kingdoms, but the combination in one molecule is unique to plants (Capella *et al*., 2016). The HD-Zip family is divided into four subfamilies (I to IV). HaHB4 and HaHB11 are divergent members belonging to sunflower, a species adapted to rather extreme climates. The ectopic expression of HaHB4 in Arabidopsis, wheat, and soybean resulted in plants with higher yield and increased tolerance to water deficit (Gonzalez *et al*., 2019, 2020; Ribichich *et al*., 2020). HaHB11 was shown to confer flooding and defoliation tolerance to maize (Raineri *et al*., 2022), and increased yield, independently of the heterosis extent (Raineri *et al*., 2019). Moreover, rice plants expressing this sunflower gene exhibit improved yield mainly due to architecture changes (Raineri *et al*., 2023). In these cases, proofs of concept were carried out in Arabidopsis plants (Dezar *et al*., 2005; Cabello *et al*., 2016), and later the transgenes were introduced in each crop and tested in greenhouses and field trials to validate the technology.

In this work, we describe the effect of the ectopic and constitutive expression of the sunflower HD-Zip I *HaHB11* in soybean plants. We obtained transgenic soybean plants using a *35S:HaHB11* construct and thoroughly evaluated the performance of two independent events over four contrasting seasons in field trials. We measured multiple parameters during crop development and observed several, albeit not all, common characteristics shared with the previously described HaHB4 soybean (Ribichich *et al*., 2020). We evaluated morphological traits, biomass, life cycle, branches and pods architecture, flowering dynamics, pollen germination under heat stress and carbohydrate distribution. Furthermore, we conducted reflectance measurements to differentiate HaHB11 plants from controls by using non-destructive methods. We concluded that HaHB11 plants achieved higher seed sets and filling than their controls due to more synchronized flowering and better pollen germination.

## Materials and methods

### Genetic constructs

The open reading frame of cDNA encoding full-length HaHB11 cloned in the *Bam*HI/*Sac*I sites of pBluescript SK-(Stratagene, Upsala, Sweden) was cloned in expression cassettes bearing the constitutive 35S CaMV promoter as previously described in a vector carrying the bar gene and the NOS termination sequence (Raineri *et al*., 2023). Clones were obtained in *Escherichia coli*, and then *Agrobacterium tumefaciens* (strain EHA101) was transformed.

### RNA isolation and expression analyses by real-time RT-PCR

Total RNA for real-time RT-PCR was isolated from soybean leaves using Trizol® reagent (Invitrogen, Carlsbad, CA, USA) and real-time qPCR was performed using an Mx3000P Multiplex qPCR system (Stratagene, La Jolla, CA, USA) as described before (Raineri *et al*., 2019). Primers used were: primer F: 5‘-ggAggAgCTAATTAAggATgTAACg-3’; primer R 5‘-gCATAgTgCAAACAAACAgCAAgC-3’ for HaHB11, and 5‘-ATggAAAAgATTTggCATCATACCTN-3’; primer R 5‘-CAgTTgTACgACCACTTgCATACAgN-3’ for internal control *ACTIN*.

### Plant transformation and selection of transgenic events

Soybean transgenic events were generated using an Agrobacterium-mediated protocol and the cultivar Williams 82 (hereafter W82) according to the methods described by Hofgen and Willmitzer (1988). Transgenic events were selected using ammonium glufosinate. T1 seeds were obtained for two independent events.

Seed multiplication was conducted in a greenhouse. T1 individuals derived from each event were sampled for a segregation test by PCR determination. Lines derived from selfings of individuals from selected events (3:1 segregation in T1) were sown and analyzed by PCR to identify homozygous lines, as indicated by the absence of negative segregants among the sampled progeny (at least five individuals sampled per line). Seed augmentation (T3 seed) of single-copy homozygous was carried out in a greenhouse. Transgenic lines bearing the HaHB11 cDNA were named H11-1 and H11-2.

### Field assays

Field assays were carried out at the IAL site on a sandy soil of 2 m depth with low water-holding capacity and an organic ‘A’ horizon of 15 cm. Evaluated germplasm included controls (null segregants and Williams 82 wild-type plants) and transgenic lines (H11-1 and H11-2). The sowing date was between November 26^th^ and November 29^th^ (depending on the campaign), using a single stand density of 30 plants m^-2^. The seeds were inoculated with *Bradyrhizobium japonicum* before sowing. The genotypes were distributed in a completely randomized design with at leastthree replicates. Each plot had three or four rows of 1.25 m and 0.3 m between rows. The plots were drip-irrigated along the whole cycle to keep the uppermost 1 m layer at field capacity and were fertilized with P (130 kg ha^-1^ at sowing). The plots were kept free of weeds, insects, and diseases. Daily maximum temperature (in °C) and incident solar radiation (in MJ m^-2^ day^-1^) were obtained from NASA POWER (https://power.larc.nasa.gov/data-access-viewer/). Heat Stress indices (HIS) was computed as the cumulated temperature above 35 °C in the evaluated period. All experiments were carried out after getting the corresponding authorization from the CONABIA (National Committee of Biotechnology) and INASE (Seeds National Institute).

### Evaluation of crop phenotype

Days from R1 to R7 were assessed in four experiments. In all assays, grain yield (GY) was obtained at R8 by hand, harvesting all the plants in at least 0.5 m of a central row of each plot, which were threshed for seed recovery.

The grain number (GN) and individual grain weight (GW) were assessed in all experiments. For this purpose, at least three samples of 100 seeds each were taken from the seed bulk and weighed; the obtained values were averaged to estimate individual seed weight. The GN was computed as the ratio between GY and individual GW and expressed on a per plant basis.

Morphological and physiological characteristics, including plant height, stem diameter, node number, node number/cm, and leaf number were recorded during the plant cycle from sowing to harvest. For this purpose, at least three plants from each plot were collected (around 12 plants per genotype). The aerial vegetative material was divided into stem (including petioles), leaf, and pods, then dried at 60 °C until constant weight. The number of flowers (divided into buds, white and senescent flowers) and pods (divided into pods smaller than 1 cm, between 1 and 5 cm, and larger than 5 cm) were counted from R1 to harvest during the 2021-2022 campaign. Pods smaller than 1 cm, between 1 and 5 cm, and larger than 5 cm were counted.

Biomass partitioning to reproductive organs was estimated as the ratio between pod weight and total biomass per plant, and between seed yield and total biomass per plant (harvest index, HI).

The light interception efficiency (quotient between intercepted and incident photosynthetically active radiation) was measured with a ceptometer (Cavadevices, Argentina) as previously described (Maddonni and Otegui, 1996).

The accumulated biomass during grain filling was computed as the difference between aerial biomass at harvest and R5.

The proportion of pods with 1, 2, 3, or 4 seeds was quantified by counting at least eight plants per genotype. The number of pods per node was counted along the stem in at least 17 plants per genotype. Node #1 was considered the first node (from the base of the plant) carrying at least one pod.

Nodulation assay was performed as previously described Song et al, 2021. Briefly, 50 seeds were surface-treated with 2 ml of *Bradyrhizobium spp.* (OD 0,5) 15 minutes before sowing in sterile vermiculite. Plants from all genotypes were grown for one month and irrigated with Fahraeus medium. Nodules were removed from the roots, photographed, and oven-dried until constant weight. The images were processed using the free ImageJ software.

### Carbohydrate, protein, and oil contents

Starch, sucrose, glucose, and protein contents from leaves, petioles and stems of at least four plants were assessed as previously described (Cabello *et al*., 2016; Raineri *et al*., 2022).

Seed composition was determined by an external service (Cámara Arbitral de Cereales de la Bolsa de Comercio de Santa Fe, https://www.cac.bcr.com.ar/es). Oil and protein contents were quantified by NIR spectroscopy.

### Pollen germination and microscopic analyses

Pollen germination and tube length were evaluated according to Salem *et al*. (2007). Briefly, recently opened soybean white flowers were harvested between 9:00 and 12:00 am on sunny days. At least 12 flowers from each genotype were dissected. Anthers were exposed, and pollen was dusted on the germination media pre-incubated at 30 °C or 45 °C in a humidity chamber by using a nylon brush (Salem *et al*., 2007). After 24 h of incubation, pollen germination and pollen tube length were assessed. Pollen grain was considered germinated when the tube length was equal to the grain diameter.

After aniline blue staining microscopic visualization was performed using an Eclipse E200 Microscope (Nikon, Tokyo, Japan) and photographed with a Nikon Coolpix L810 camera. The tube length was measured using the ImageJ software.

### Remote sensing analysis

Canopy spectral reflectance was measured using a compact shortwave NIR spectrometer (Ocean Insight) as previously described for maize plants (Raineri et al., 2022).

Measurements were collected on 01/17/2020 (flowering), 01/28/2020 (pod development), and 03/02/2020 (late seed development). A typical outlier control based on standard deviation was implemented on each canopy spectral reflectance raw data. Twenty-six spectral vegetation indices (VIs) were selected based on the range of available wavelengths. Such VIs were related to biomass accumulation, plant stress conditions, and chlorophyll content. We also included a temperature-based index: canopy-air temperature differential (Tc – Ta), in which Tc is the canopy temperature and Ta, is the air temperature.

The selected indices, their formulas, and applications are shown in Supplementary Fig. S7 and Table S2.

Each vegetation index was evaluated for the controls and the two transgenic lines. Data analyses were conducted in R using the aov function, and the post-hoc test was performed using the agricolae-package (Mendiburu, 2010).

### Statistical analyses

Differences in seed yield and its components, pods/plant, pods/node, individual pod weight, total biomass and harvest index between WT cv. W82 and TG events and between W82 and TG cv. H11-1 and H11-2 were assessed by means of analyses of variance (ANOVAs) using Infostat/L software, with genotypes (G), and event (E) as classification factors and campaign (C) as fixed factor. The model used was genotype > event, genotype > campaign, event > campaign. An LSD Fisher test was used for comparison. Significant effects represented p value <0.1 (*); 0.05 (**), or 0.01 (***), respectively. Principal component analysis (PCA) was conducted using Infostat/L software. The analysis included the variables depicted in Fig. 3, with campaign, genotype, and event serving as classification variables. The resulting correlation matrix and biplot are shown in Fig. 3.

## Accession numbers

For sunflower HaHB11 and HaHB4, accession numbers in EMBL, GenBank and DDBJ Nucleotide Sequence Databases are Ha412T4l19912C0S1, and AF339748 and AF339749, respectively.

## Results

### Soybean plants expressing the sunflower transcription factor HaHB11 differ from controls in the number of nodes, leaves, and total biomass at specific developmental stages

The sunflower *HaHB11* gene was shown to confer beneficial characteristics to Arabidopsis, maize, and rice plants. Such traits differed between the transformed species, except for one shared by all of them: grain yield (Cabello *et al*., 2016, Raineri *et al*., 2019, 2022, 2023). To test if the sunflower gene can improve soybean yield, we transformed the Williams 82 variety with a *35S:HaHB11* construct and obtained two independent events named H11-1 and H11-2. Expression levels of the transgene in those events, evaluated by RT-qPCR, were similar (Fig. 1A). Since there is not an endogenous homolog, we could only compare such expression with that obtained in other transgenic crops, being similar. These genotypes were evaluated in four field trials conducted in 2019, 2020, 2021, and 2022 in micro plots under irrigated conditions.

**Figure 1:**
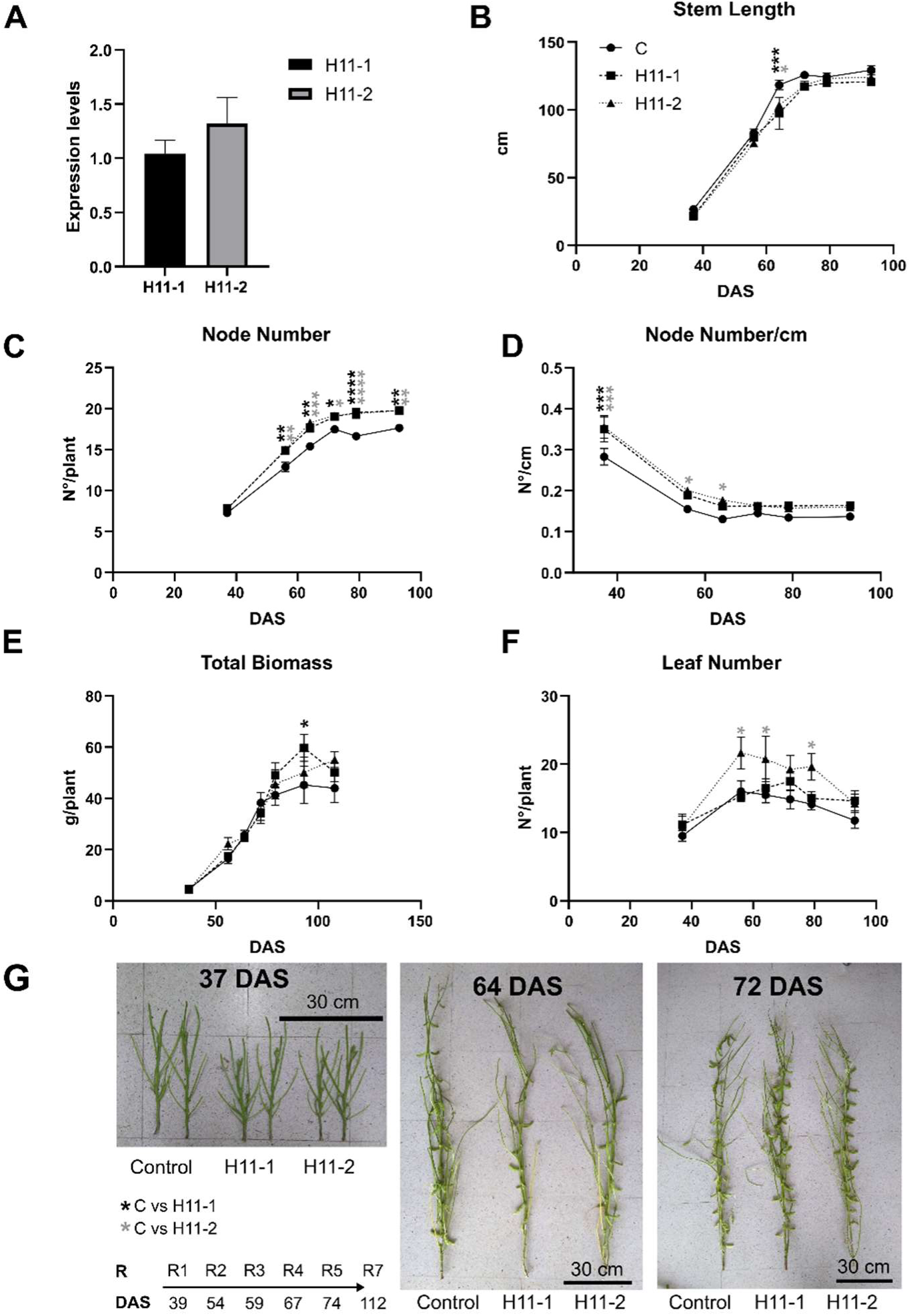
Soybean plants expressing the sunflower *HaHB11* gene exhibit more nodes, leaves, and biomass than their controls at specific developmental stages. (A): *HaHB11* expression levels in two independent events (H11-1 and H11-2). (B): stem length; (C): node Number/plant, (D): node number/cm; (E): total biomass, and (F): leaf number during development. (G): Illustrative images of control, H11-1, and H11-2 events at 37, 64, and 72 days after sowing (DAS). In the bottom left: days required to arrive developmental stages R1 to R7. Data were analyzed using a two-way ANOVA considering genotype and treatment, followed by a Tukey test. Black and grey asterisks (*) indicate significant differences (* p<0.05; **p<0.001; ***p<0.0001) between C and H11-1, and C and H11-2 plants, respectively. The showed data corresponds to the 2019-2020 campaign.

The mean plant growth characterization of each event across environments is shown in Fig. 1. Stem elongation tended to be slower in H11 plants than in controls, particularly between R1 and R4, being those of H11 the last to reach maximum plant height approximately at R5 (Fig. 1B and Supplementary Fig. S1A). Node number per plant was always higher in both transgenic events from R3 onwards (Fig. 1C), slightly varying among growing seasons (Supplementary Fig. S1B). Node density (in nodes per cm) was consistently higher in both transgenic events up to 60 days after sowing at the R3 stage (Fig. 1D and Supplementary Fig. S1C). Despite these differences, ground cover evolution was similar between H11 and control plants (Supplementary Fig. S1F). Considering nodulation, no significant differences were detected among genotypes in the number of nodules per plant or their volume (Supplementary Fig. 1G-I).

Total biomass and leaf number per plant also differed between control and transgenic plants, depending on the developmental stage (Fig. 1E-1F; Supplementary Fig. S1D-S1E). Regarding the growing rate, although H11 plants were shorter on 34 days after sowing (DAS), they became similar in height on 64 DAS and taller on 72 DAS (Fig. 1G). Trends for these traits were similar across growing seasons (Supplementary Fig. S1), indicating a strong consistency in the effect of the sunflower transgene.

### H11 plants exhibit an increased grain number compared to controls

Together with the previously described characteristics evaluated along the vegetative and reproductive stages, we were particularly interested in analyzing traits affecting grain set and filling. Grain yield and its components were assessed in the four growing seasons, differing slightly across them (Table 1). Transgenic plants bearing the sunflower HaHB4 gene were added in the last season as a reference genotype, given that such plants had been well-characterized in previous research (Ribichich *et al*., 2020). Although not statistically significant, transgenic plants always outyielded the controls in three out of four trials. The mean increment in grain yield across growing seasons was 10%. Grain number, grain weight, biomass production, and harvest index were evaluated to understand which components led to the observed trend in grain yield.

**Table 1.** H11 plants outyielded controls in three out of four campaigns. Traits assessed during four field trials carried out between 2018 and 2022. Values and standard deviation of the media (SEM) are shown for two independent events of soybean plants transformed with HaHB11 (H11-1 and H11-2). Evaluation of soybean plants transformed with HaHB4 (H4) is shown for the campaign 2021-2022.

|  | 2018-2019 | 2019-2020 | 2020-2021 | 2021-2022 |
| --- | --- | --- | --- | --- |
| **Yield/ plant (g/plant)** |  |  |  |  |
| **CONTROL** | 10 ± 0.9 | 14 ± 1.7 | 16 ± 0.8 | 19 ± 1.3 |
| **H11-1** | 12 ± 1.0 | 16 ± 1.5 | 15 ± 0.8 | 20 ± 1.1 |
| **H11-2** | 11 ± 0.7 | 15 ± 1.2 | 16 ± 1.2 | 21 ± 1.2 |
| **H4** |  |  |  | 19 ± 1.0 |
| **Grain N°/plant (total N°/plant)** |  |  |  |  |
| **CONTROL** | 53 ± 5.3 | 70 ± 7.9 | 75 ± 4.1 | 82 ± 4.7 |
| **H11-1** | 66 ± 3.7 | 82 ± 7.6 | 80 ± 7.7 | 102 ± 4.8 |
| **H11-2** | 69 ± 3.8 | 85 ± 6.2 | 79 ± 6.5 | 111 ± 5.8 |
| **H4** |  |  |  | 99 ± 5.3 |
| **100 grains weight (g/100 grains)** |  |  |  |  |
| **CONTROL** | 20 ± 0.8 | 19 ± 0.4 | 22 ± 1.1 | 22 ± 0.0 |
| **H11-1** | 17 ± 1.9 | 19 ± 0.5 | 20 ± 0.5 | 22 ± 0.3 |
| **H11-2** | 16 ± 0.7 | 17 ± 0.5 | 19 ± 1.0 | 19 ± 0.2 |
| **H4** |  |  |  | 19 ± 0.3 |
| **HI (grain weight/total biomass)** |  |  |  |  |
| **CONTROL** | 0.24 | 0.31 | 0.308 | 0.328 |
| **H11-1** | 0.28 | 0.26 | 0.280 | 0.407 |
| **H11-2** | 0.29 | 0.30 | 0.296 | 0.392 |
| **H4** |  |  |  | 0.327 |

Table 2 summarizes the study performed with the three genotypes (H11-1, H11-2, and controls Williams 82). H11 plants exhibited an increased grain number compared to controls. Despite their lighter pods and grains, they developed more total plant biomass than controls, a higher number of nodes and pods per plant, and pods per node, resulting in an increased number of grains per plant. Although H11 plants fixed more grains, yield differences were not statistically significant (Table 2), probably due to a trend to penalize harvest index in some environments. As mentioned above, the picture was different, considering each trial individually (Table 1).

**Table 2:** H11 plants significantly differ from controls in multiple traits across four field trials. The table summarizes data variance analysis considering genotype and treatment. The traits shown in the table were assessed during four campaigns. An ANOVA analysis followed by an LSD Fisher test was performed. The model considers the following hierarchies: genotype>event, genotype>campaign, event>campaign. H11 comprises both independent events. All the values were compared with those measured in control plants. Those highlighted in green represent increased values, whereas those in yellow, decreased ones. Black and grey asterisks (*) indicate significant differences (* p<0.05; **p<0.001; ***p<0.0001) between C and H11-1 and C and H11-2 plants, respectively.

| Genotype<br>(compared<br>with control) | Grain number | Pods/<br>plant | Nodes<br>/<br>plant | Pods/<br>node | Grain<br>weight | Pod<br>weight | Grain<br>yield | Total<br>biomass | HI |
| --- | --- | --- | --- | --- | --- | --- | --- | --- | --- |
| H11 | *** | *** | *** | *** | *** | *** | - | *** | - |
| Event<br>(compared<br>with control) |  |  |  |  |  |  |  |  |  |
| H11-1 | *** | *** | ** | ** | - | *** | - | * | * |
| H11-2 | *** | *** | ** | ** | *** | *** | - | - | - |
| Up | Down |  |  |  |  |  |  |  |  |

On the one hand, grain yield was strongly and positively correlated with grain number (r = 0.92; p < 0.001), whereas the correlation with individual grain weight was weaker (p = 0.05, Supplementary Fig. S2). The grain number was significantly larger for H11 than for controls (Tables 1 and 2). The only season in which H11 plants did not outyield controls was that in which grain number did not show differences among genotypes. For all the traits considered, the expression of *HaHB11* brought an increase in grain number and a decrease in grain weight, indicating that the trend to improved seed yield conferred by *HaHB11* was driven by a marked improvement in the first component, compensated by a decline in the second one. Physiological steps conducive to this trade-off between grain yield components are analyzed next.

### H11 plants develop more flowers per node than their controls

To unravel the origin of grain number differences, we assessed traits conducive to its determination, particularly focusing on flower developmental kinetics and pod fixation. Both H11 events bloomed only 1-3 days later than their controls and achieved a higher number of flowers per plant and per fertile node between R2 and R4 (Fig. 2A-2B; Supplementary Fig. S3). Buds, newly opened flowers, and senescent flowers (which summed up constitute the total number of flowers) were counted, showing similar trends to those described in Fig. 2 (Supplementary Fig. S3). Moreover, the number of set pods per node was significantly higher in the transgenic events compared with controls (Table 2, Fig. 2C), and notably concentrated mainly between nodes 4 and 6 in the mid-lower section of the plants (Fig. 2D), although the initial developmental rate (pods/node) was slightly lower than those of controls (Fig. 2C and Supplementary Fig. S4). Pods per plant were also classified into different sizes: pods with less than 1 cm, between 1 and 5 cm, and larger than 5 cm. No significant differences were observed in these traits (Fig. S3). To understand whether the difference in grain number was related to the number of seeds per pod, we estimated the proportion of pods with 1, 2, 3, or 4 grains per pod. There was no difference between genotypes (Supplementary Fig. S4E). Protein and oil contents were evaluated in the three genotypes to test grain quality, showing a similar composition (Supplementary Fig. S4F).

**Figure 2:**
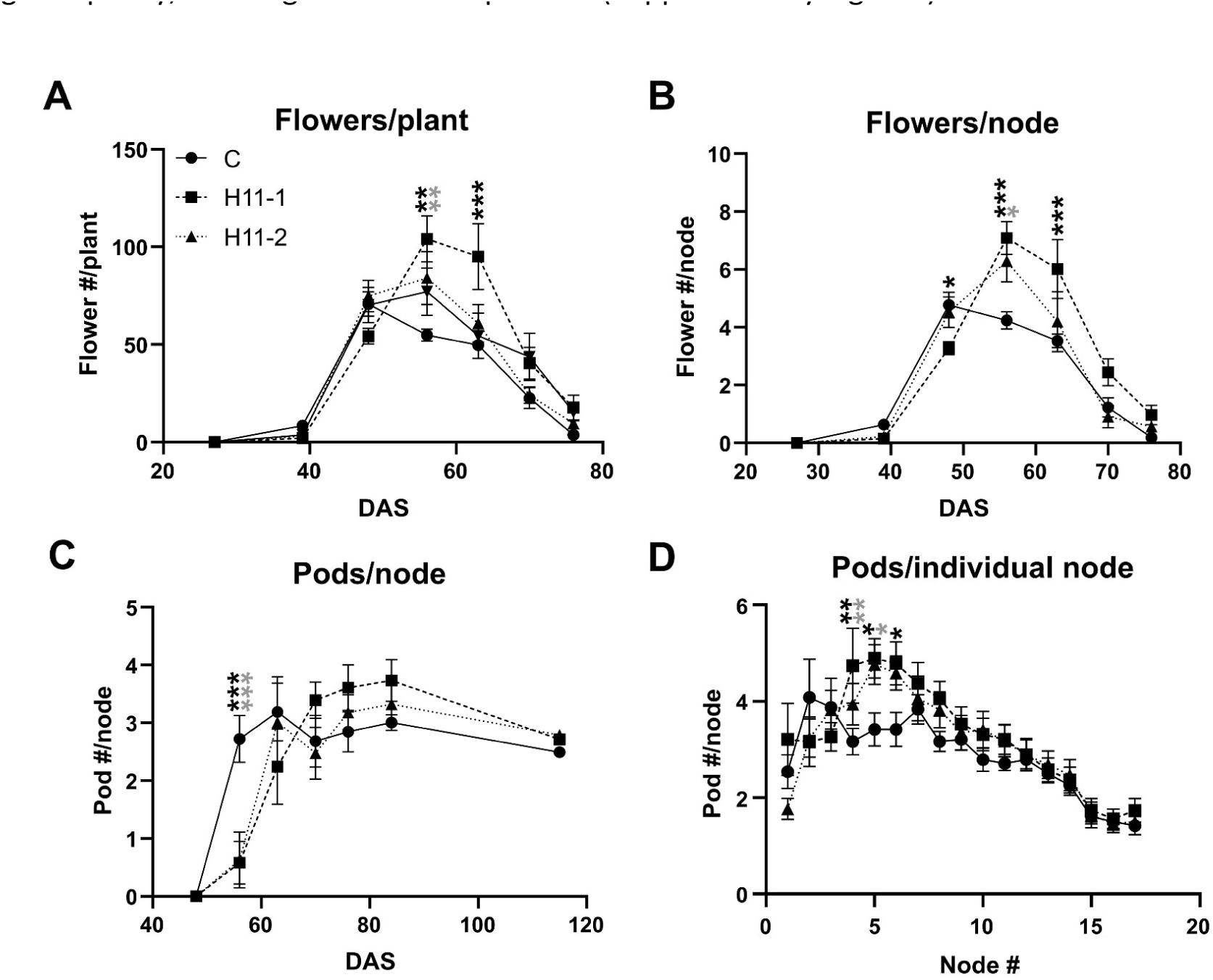
H11 plants have more flowers per plant than their controls. (A): flower number per plant, (B): flower number per node, and (C): pod number per node. (D): pod number per node from node 1 (basal) to node 19 (apical). At the bottom (left), the number of days required to reach the developmental stages from R1 to R7 is indicated. DAS: days after sowing. Data were analyzed using a two-way ANOVA followed by a Tukey test. Black and grey asterisks (*) indicate significant differences (* p<0.05; **p<0.001; ***p<0.0001) between C and H11-1 and C and H11-2 plants, respectively.

### The increased grain number observed in H11 plants could be associated with improved reproductive performance under high temperature

Given that increased grain number was detected as a robust differential trait in H11 plants, the question we asked was why this difference did not always impact grain yield positively and the effect depended upon the campaign (i.e. the environment) due to a variable trade-off effect with grain weight. Considering that the genotype characteristics are stable, the hypothesis was around the effect of environmental conditions. We conducted a comprehensive principal component analysis (PCA) to reveal a potential mechanism involved in the differential phenotype. Such analysis included the physiological determinants of grain yield (i.e. total shoot biomass and harvest index), its components (grain number and individual grain weight) and growing conditions during the flowering period (R1-R3), the grain set period (R3-R5) and the grain-filling period (R5 onwards). Since the four trials were uniformly irrigated and conducted on similar sowing dates, temperature and irradiance emerged as pivotal factors influencing grain number. For this reason, we included the Average Maximum Temperature (Tm, in °C) as a discriminant environmental variable during the specified growth stages, considering it is a better indicator of differences across years compared to the daily mean during the summer period. We also added Accumulated Incident Irradiance, because it is the main driving factor of biomass production differences in an irrigated environment (Fig. 3A).

**Figure 3:**
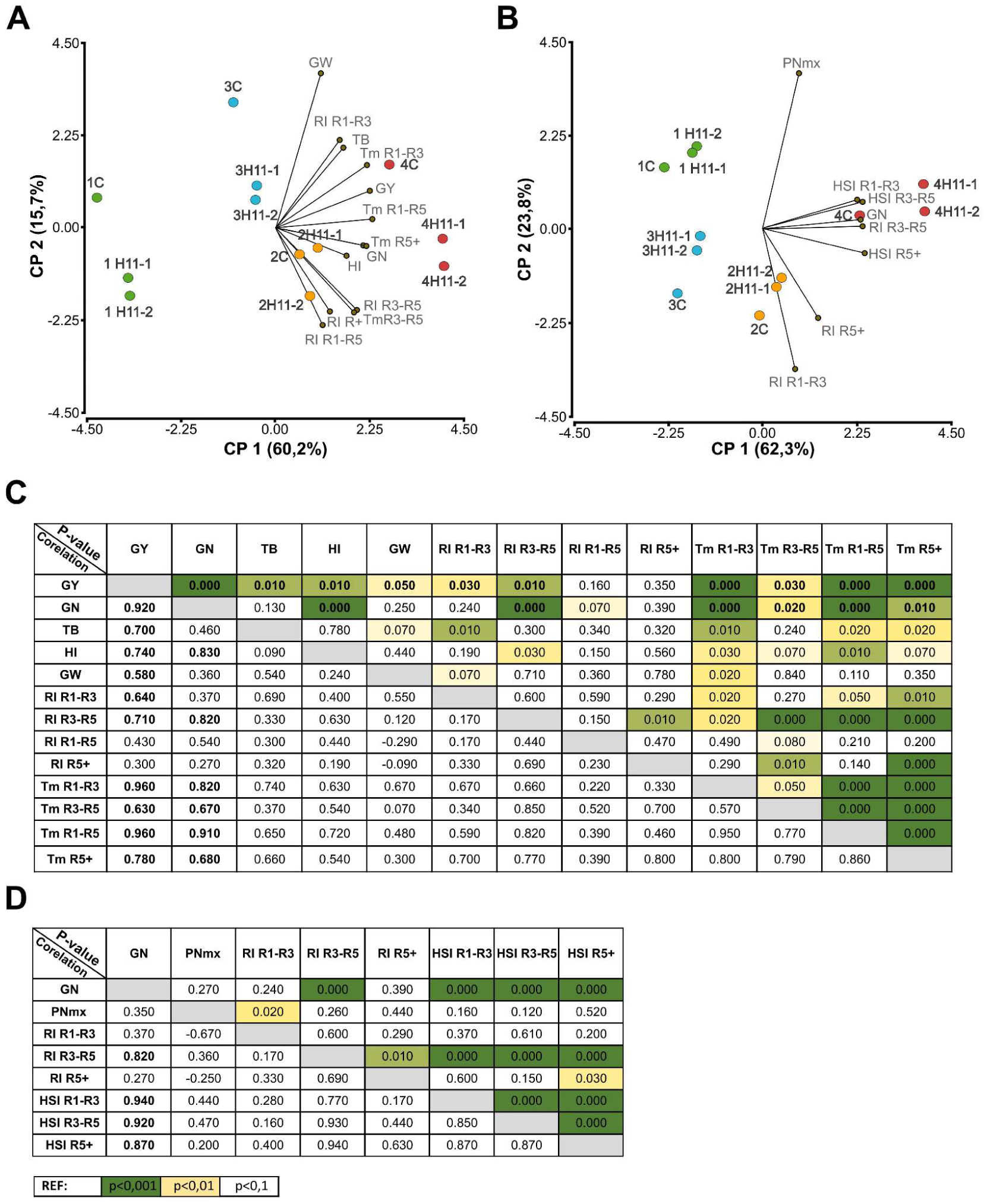
Grain number, grain yield, and total biomass positively correlate with the temperature registered between R1-R2. PCA analysis includes four field trials with control, H11-1, and H11-2 plants. The biplot (A and B), correlation, and P-value matrix (D). P-values are indicated using different colors to represent significant levels of 0.001 (green), 0.01 (yellow), and 0.1 (white). GN (Grain Number/plant); Y (Yield); TB (Total Biomass/plant); HI (Harvest Index); GW (Grain Weight); PNmx (Maximum number of pods/plant). Average Maximum Temperature (Tm), Cumulative Incident Irradiance (RI), and Heat stress index (HSI) were computed between R1-R3, R1-R5, R3-R5 and R5 to the end of cycle (R5+).

Grain yield was significantly correlated (p = 0.01; Fig. 3C) with both physiological determinants (total shoot biomass and harvest index) and both numerical components (p ≤ 0.05; Fig. 3C). The relationship was strongest with grain number (p < 0.001; Fig. 3C). The PCA explained 75.9 % of the joint variation among all these traits and the selected environmental variables (Fig. 3A), with 60.2 % accommodated in the PC1 and 15.7 % in the PC1. On the one hand, the major part of the variation registered in grain yield, grain number and harvest index were encompassed by the first component, together with mean maximum temperatures along the cycle. Comparatively higher values of these traits deployed towards positive values of the *x* axis and smaller values towards the negative side of this axis. On the other hand, almost all the variation registered in individual grain weight was encompassed by the second component, with comparatively larger values of the trait towards positive values of the *y* axis and vice versa. The major part of the variation in cumulative incident irradiance along the cycle (R1-R5 and R5 onwards) was also represented along the PC2 but with comparatively larger values towards negative values of the *y* axis. The variation in total aerial biomass was distributed between the positive trajectory of both axes, a trend accompanied by cumulative irradiance during the flowering period (R1-R3) but not by cumulative irradiance and mean maximum temperature during the pod set period (R3-R5), which increased towards positive values of the *x* axis and negative values of the *y* axis (Fig. 3A).

The marked effect of the environment on grain yield and its main component (grain number) was evident in the distribution of experiments along the PC1, which clearly distinguished the growing season with the highest grain yield (2021-2022) on the right extreme of the *x* axis and the growing season with the lowest grain yield (2018-2019) on the left extreme of it (Fig. 3A). The other seasons, with intermediate grain yields, were located close to the center of the PC1 (*x* = 0). However, season two (2019-2020) data deployed towards positive *x* values and negative *y* values, indicative of its grain yield defined by a higher number of grains of smaller weight with respect to season three (2021-2022) that showed the opposite trend (Fig. 3A). Within each season, H11 genotypes always outperformed the controls in grain number, a trend that usually translated into improved grain yield of the former. The exception to this trend occurred in season three because of a complete trade-off between the components (the perpendicular projection of the data corresponding to the Control intercepted the grain weight vector at the most distant positive point from the origin).

Additionally, it is important to highlight that no association was detected between the physiological determinants of grain yield or between its components (as indicated by vectors at nearly right angles). This pattern allows selection for one trait without compensation by the other (Fig. 3A, as indicated by vectors at obtuse angles), provided both traits are included in the selection process. Considering the strong relationship between grain yield and grain number and the relevance of high temperature all along the cycle on the determination of the latter, we developed a new PCA including a heat stress index (HSI; Fig. 3B-D) to explore the impact of this condition on grain number determination and the improved relative performance of H11 respect to controls for this trait. The HSI represented the cumulative degrees above 35 °C on a daily basis (HIS= Tm – 35, for Tm > 35 °C) and was computed for each variety × growth stage × growing season combination (Supplementary Table S1). The analysis included the maximum number of pods per plant (PNmx) as a proxy of potential sink size. The principal components accommodated 86.1 % of the variation corresponding to the selected variables, 62.3 % by the PC1 and 23.8 % by the PC2 (Figure 3B). Almost all the variation registered in grain number was displayed along the first component, with larger relative values towards the right of the *x* axis. This trend was strongly accompanied (sharp acute angles) by all evaluated environmental variables except the amount of incident irradiance early (RI R1-R3) and late (RI R5+) in the cycle (vector in right angle denoting no association). The difference in grain number favoring H11 genotypes was largest in the environment with the highest HSI during R1-R3 and R3-R5 (2021-2022 Supplementary Table S1, and Fig. 3). The second component accommodated the variation registered in the maximum number of pods per plant (towards positive values of the *y* axis) and in cumulative daily irradiance early (RI R1-R3) and late (RI R5+) in the cycle (towards negative values of the *y* axis). All these variables were at the right angle with the grain number vector, indicative of no relationship with it (Fig. 3B-3D).

### The pollen from H11 and H4 plants germinate better than that of controls at high temperatures

Considering the PCA analysis shown above, we aimed to understand why H11 plants fixed more grains than controls at higher mean maximum temperatures. Pollen germination and tube growth are susceptible to heat stress and suppressed at temperatures above 50 °C. Hence, we decided to compare H11 and control pollen germination under heat stress (Fig. 4). While pollen from all genotypes fully germinated at 30 °C, less than 10 % of pollen from control plants germinated at 45°C, whereas that of H11 flowers showed a germination rate between 35 and 60 % (Fig. 4). Transgenic soybeans expressing *HaHB4* were shown to be tolerant to heat stress across 27 field trials (Ribichich *et al*., 2020). Notably, although other traits were clearly different between HB4 and H11 soybeans, both genotypes display enhanced grain settings (Ribichich *et al*., 2020) with a minimum delay in flowering. We evaluated HB4 pollen germination at 45° and found similar results to those of HB11. HB4 pollen germination was around 3-fold higher than that of controls. Additionally, H11 and HB4 pollen grains tended to develop longer pollen tubes under heat stress compared to controls (Fig. 4).

**Figure 4:**
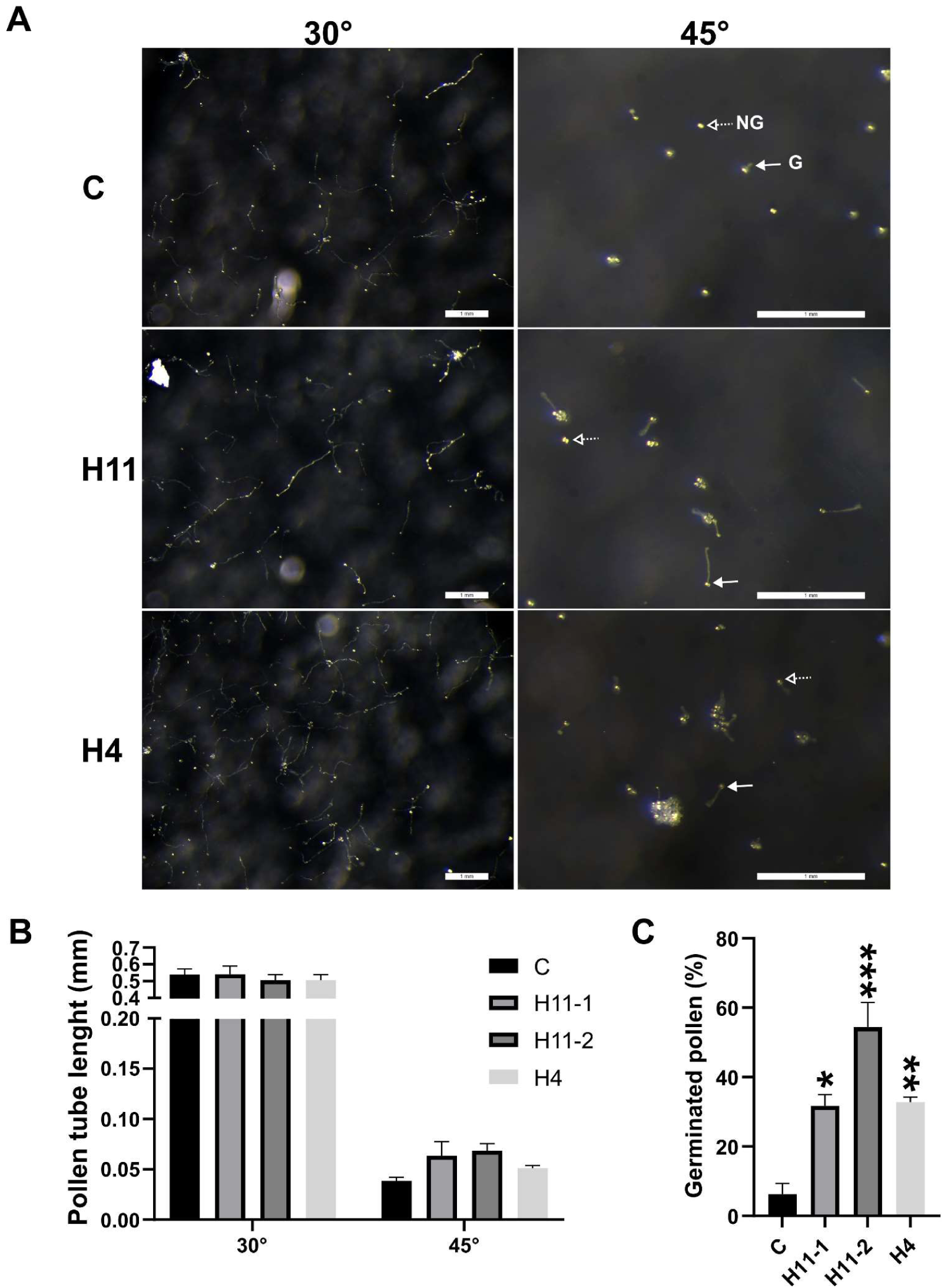
H4 and H11 pollen grains germinate better than controls under heat stress. (A): Illustrative pictures of pollen grains of control (C), transgenic *HaHB11* (H11), and transgenic *HaHB4* (H4) plants at 30°C and 45°C, respectively. Dashed-open arrows point not germinated pollen (NG), and solid ones, to germinated pollen (G). The pictures were taken with a microscope; white bars indicate 1 mm. (B): Quantification of pollen tube length at 30°C (normal conditions) and 45°C (heat stress). (C): percentage of germinated pollen at 45° for each genotype. Data were analyzed using a two-way ANOVA considering genotype and treatment followed by a Tukey test. Asterisks indicate significant differences (* p<0.05; **p<0.001; ***p<0.0001).

### H11 and HB4 plants differentially distribute their carbohydrates throughout the plant

The improved pollen grain germination under heat stress exhibited by the transgenic plants can explain the observed higher number of pollinated flowers. However, to set more pods and grains, it is necessary to have more nutrients. To investigate this issue, we evaluated the carbohydrate (glucose, sucrose, and starch) and protein contents at the R5 stage, when grain filling begins. We sampled stems, petioles, and leaves from nodes 5, 8, and 15 (lower, middle, and upper parts, respectively) of plants to gain insight into the distribution of these biomolecules (Fig. 5).

**Figure 5:**
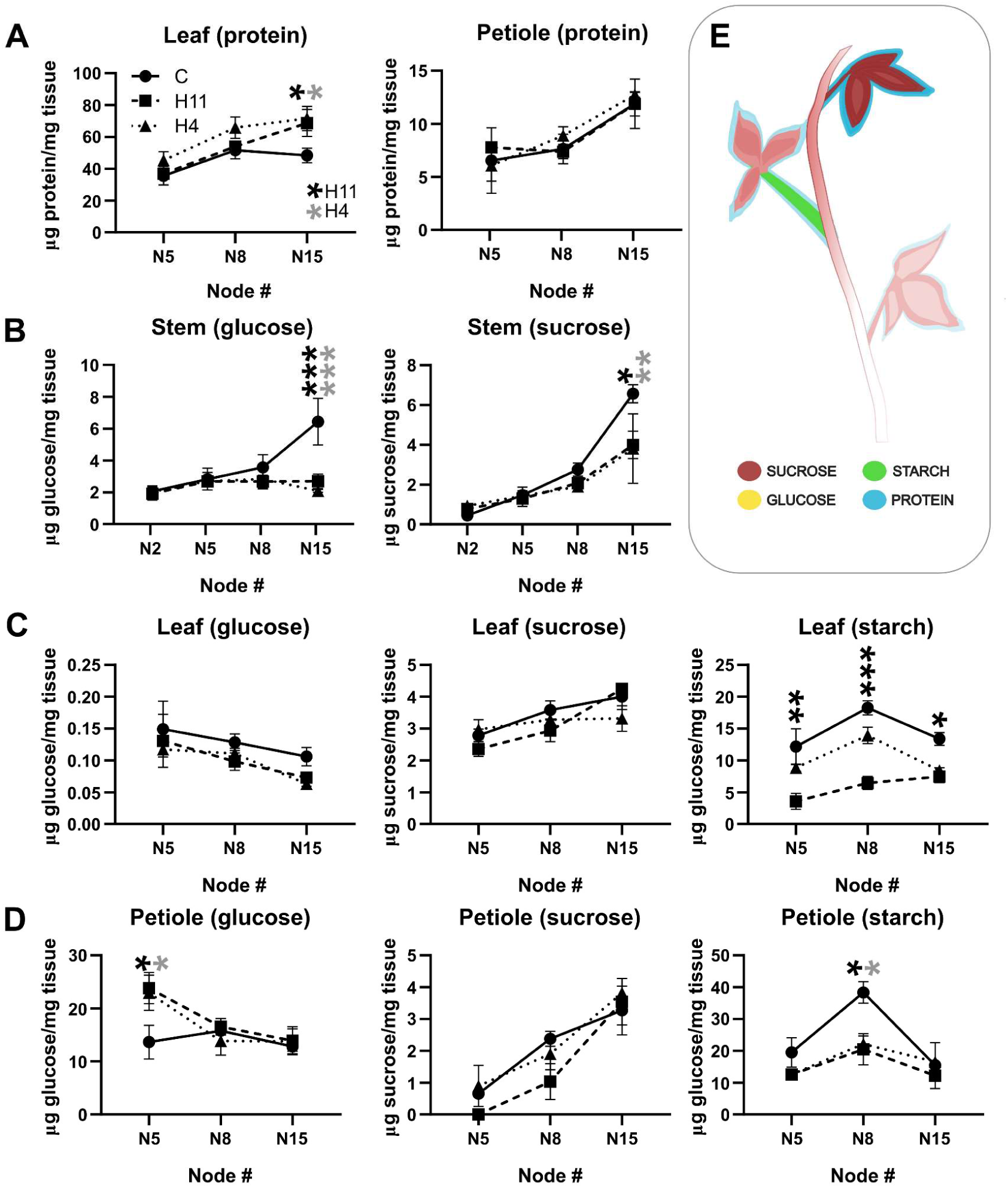
H11 plants display carbohydrate depletion compared to their control counterparts. (A): protein content in nodes 5, 8, and 15 (N5, N8, and N15) of leaves and petioles. (B): glucose and sucrose contents in nodes 2, 5, 8, and 15 of the stem. (C) and (D): glucose, sucrose, and starch contents in nodes 5, 8, and 15 of leaves (C) and petioles (D). Schematic representation showing the carbohydrate and protein contents of Williams 82 soybean plants at the R5 stage in the evaluated tissues (E). Data were analyzed using a two-way ANOVA considering genotype and treatment followed by a Tukey test. Black and grey asterisks (*) indicate significant differences (* p<0.05; **p<0.001; ***p<0.0001) between C and H11 and C and H4 plants, respectively.

H11 and HB4 plants showed lower sucrose and glucose contents and higher protein content at the apex of the stem than the controls. Moreover, they had less glucose and starch contents in their leaves. In middle and basal petioles, starch content was lower than in controls and glucose higher (Fig. 5). Considering the relationship between the accumulated biomass during grain filling (R5 to the end of the cycle) and the individual grain weight, H11 plants were less dependent on biomass availability than controls (Supplementary Fig. S6C).

### Vegetation indexes can differentiate H11 from control plants

Proximal sensing offers a rapid and non-destructive approach to plant screening, allowing to account for environmental factors and also improving the accuracy of quantifying plant phenology development. Reflectance measurements were taken at different developmental stages of control and H11 plants during the 2019-2020 campaign and used to generate a set of 26 spectral vegetation indices (VIs). VIs capable of discriminating between H11 and control plants were identified by statistical analysis. Welch’s t-test was employed due to assumed unequal variances between transgenic crops. During early reproductive stages (R2 and R3), six VIs discriminated between transgenics and controls (Fig. 6; Supplementary Fig. S7). The temperature-based index could discriminate genotypes for R5 and R6.

**Figure 6:**
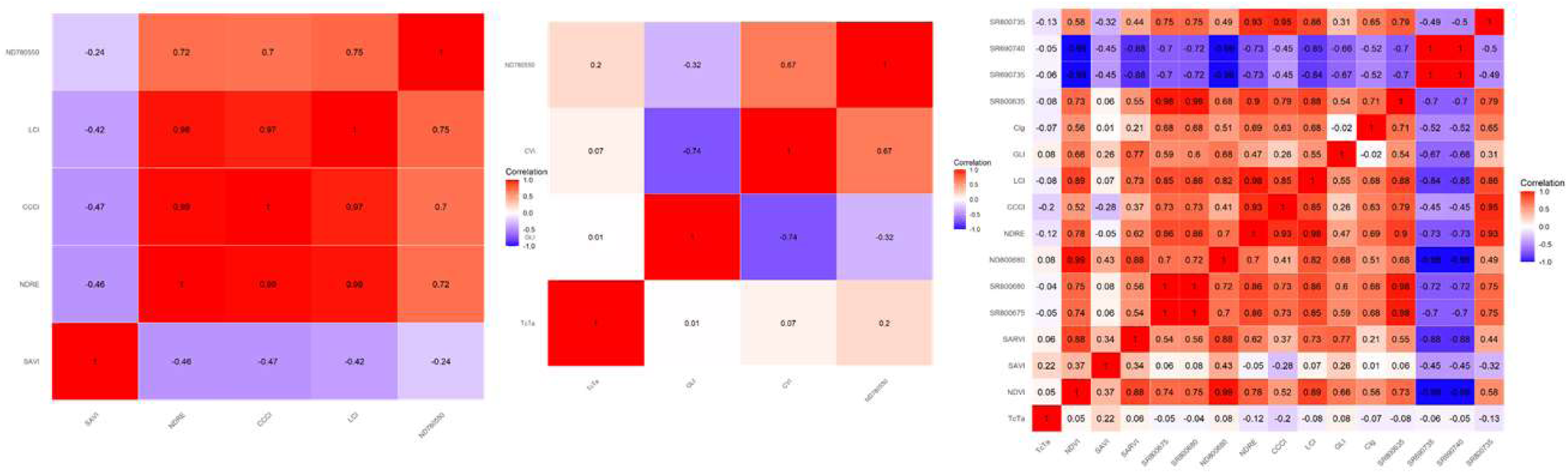
Vegetation indices computing spectral data can distinguish H11 from control plants. Correlation matrices between vegetation indices for R3, R5 and R6. Weak and moderate correlation indices for R3: SAVI; for R5: Tc-Ta, GLI, ND780-500; and for R6: Tc-Ta, SAVI, SARVI, CCCI, GLI, CIgreen, SR800-735. From left to right, second, third and fourth date of spectral sampling.

We constructed correlation matrices using Pearson’s coefficients to determine the relationships between indices capable of discriminating genotypes and to avoid redundancy. Weak and moderate correlations (below 0.60) indicated a lower association between VIs, suggesting that those indices are best suited for discrimination. Selected VI were those for which at least 50 % of interactions were weak or moderate, resulting in one VI for R3, three for R5, and seven for R6 (Fig. 6).

## Discussion

Soybean (Glycine max L. Merr.) is one of the more important crops worldwide, and it is used for human nutrition, animal feed, oil production, etc. Ninety percent of the cropped soybean is genetically modified (GM), and such modifications belong to the first transgenic generation, resistant to herbicide or herbivory attack. Despite the constant and multidisciplinary research, soybeans exhibiting abiotic stress tolerance, nutritional or quality improvement, and increased yield are absent in the market (Chan *et al*., 2020; González *et al*., 2020). We can find examples of field-evaluated GM soybeans exhibiting yield improvement. Soybean plants overexpressing the GmWRI1b transcription factor exhibited a higher seed number, containing more lipids than their controls (Guo *et al*., 2020). Seed number and weight also increased by overexpressing *GmmiR156b*, a regulator of *SQUAMOSA PROMOTER BINDING PROTEIN-LIKE*, which plays a central role in stem architecture and branching (Sun *et al*., 2019). Overexpression of the Arabidopsis gene *AtSINA2*, belonging to the RING E3 ligase family, also resulted in soybean plants with early flowering, increased plant height, enhanced drought tolerance, and augmented number of seeds and yield (Yang *et al*., 2023). Although the overexpression of all these genes, like *HaHB4* and *HaHB11* (Raineri *et al*., 2019; Ribichich *et al*., 2020), leads to a common trait, increased seed number, the molecular pathways triggered are different.

The number of fixed seeds is determined by multiple environmental and crop-inherent factors. We hypothesized that improved seed-setting of H4 and H11 was due to the increase in the number of flowers and pods per node and plant (Fig. 2; Supplementary Fig. S3-S4) even under heat stress. The experimental data corroborated the hypothesis (Table 2), in agreement with a correlation between the number of flowers and fixed pods (Egli, 2005). Despite this correlation, not all the flowers develop into pods; moreover, up to 36 % to 81 % of the developed flowers aborted (Egli, 2023). Interestingly, H4 and H11 plants significantly differed in the number of fixed pods between nodes four and six (Fig. 2). Pod fixation at each node does not strictly depend on the photosynthesis of the adjacent leaf but on the interaction between pods at the same node (Egli and Bruening, 2006). Defoliation of a node does not induce pod abortion at that node, but if defoliation occurs in the three adjacent nodes, pod fixation decreases by 50 % (Egli, 2005). More recently, the same research group observed that the “one node” model (which confines photosynthates to the node where they were generated), is a better predictor of plant behavior than the “one pool” model (which averages the entire plant, Egli, 2015). The differential distribution of carbohydrates in H4 and H11 plants (Fig. 5) could be due to average higher demand from the increased number of developing pods (sink) between nodes 4 and 6. Wild-type plants, which set fewer pods at these particular nodes (and in the whole plant), accumulated excessive reserve substances such as starch (Fig. 5C).

Synchronous flowering increases pod fixation by reducing competition between pods at the early and late stages of development (Egli and Bruening, 2002), and pods larger than 1 cm have a reduced probability of abortion (Egli and Bruening, 2006). H4 and H11 plants set pods slower than controls (Supplementary Fig. S6A), having more 1-cm pods and less 5-cm ones until 84 DAS. However, on 88 DAS the pods of all the genotypes reached their maximum and did not present differences between them (Supplementary Fig. S6). Among the transgenic events, H11-1 showed the most marked delay in the fixed pods’ growth, albeit exhibiting the highest number of pods smaller than 1 cm. H11-2 and H4 plants showed a similar, though less pronounced, trend between 70 and 76 DAS (onset of R5). The three genotypes significantly differed in their slopes of the fitted models for the pod number smaller than 1 cm, being the control the less pronounced (Supplementary Fig. S6B). This was in agreement with a more synchronous growth of the transgenic genotypes, at least for pods smaller than 1-cm, which correlates with pod and seed number (Supplementary Fig. S5).

Crops like maize, sorghum, sunflower, or soybean set a number of seeds related to the biomass accumulated during the seed set period (Andrade *et al*., 1999; Kantolic and Slafer, 2001; Vega *et al*., 2001; Rossini *et al*., 2011). In a study performed in two environments (USA and Argentina) with 86 soybean cultivars, biomass accumulation, partitioning to seeds, and seed set efficiency were evaluated, aiming to identify the prevalence of each of these traits in the seed set. The authors concluded that a higher number of seeds was related to the growth rate of the crop between R1 and R5 (seed set period) in most environments tested. However, they did not identify a sole individual trait as the main one for all the environments (Rotundo *et al*., 2012). In three of the four seasons, H11 plants outyielded controls by 5-10 % (Table 1). However, such a difference was not statistically significant when analyzing the four seasons together (Table 2). Considering biomass partitioning to pods (Supplementary Fig. S5), H11 plants tended to be less efficient than controls, especially in the 2020-2021 season. Notably, in that campaign, no yield differences between genotypes were assessed. In the campaigns in which H11 plants outyielded controls, they accumulated more biomass than controls between R3 and the end of the cycle (Supplementary Fig. S5).

The PCA for grain yield determination considered all the data for the four seasons and revealed a relationship with Tm between R1 and R5 (Fig. 3A). Further studies will be necessary to reinforced this finding, detecting a strong relationship between this grain yield component and temperatures above a threshold of 35 °C, usually representative of heat stress (Fig. 3B; Supplementary Table S1). The pollen is sensitive to high temperatures (above 32°C), leading to decreased germination and pollen tube length, regardless of maturity group (Salem *et al*., 2007; Djanaguiraman *et al*., 2013). The capability of pollen germination at high temperatures is a trait that allows the identification of soybean varieties tolerant to thermal stress (Salem *et al*., 2007; Djanaguiraman *et al*., 2019).

However, leaves and pollen tolerance to high temperatures are not necessarily related (Salem *et al*., 2007). The pollen of H11 and H4 plants germinated better than that of controls at 45°C (Fig. 4) and tended to exhibit longer pollen tubes. These data would explain why transgenic plants outyielded controls through improved grain set in our region, frequently affected by heat stress. Supporting the latter, the Tm between R3 and R5 (between 60 and 76 DAS) in the only season when H11 plants yielded similarly to controls (season 2020-2021) had the lowest temperatures of the four seasons (Supplementary Table S1). Thermal stress tolerance during the reproductive period is a crucial trait, given that global warming predicts a frequency increase of heat waves and rising temperatures (Jagadish *et al*., 2021).

Notably, in three out of four seasons, H11 plants exhibited a penalty in grain weight compared to controls, regardless of their net yield or the number of seeds they set. H11 rice and a few trials with H11 maize plants showed similar characteristics (Raineri *et al*., 2022, 2023). Soybean seeds greatly respond to source-sink particularly those that decrease the source per set grain (Borrás *et al*., 2004). Improved grain numbers are frequently accompanied by a reduction in the source per seed that usually represents a decrease source-sink ratio (Borrás *et al*., 2004). Except in one experiment, in current research increased grain number of H11 genotypes did not represent a total trade-off between both grain yield components, as observed in other species when grain set was improved by means of artificial synchronous pollination (Uribelarrea *et al*., 2008). Albeit the carbohydrate depletion in H11 plants (Fig. 5) partially explains this source restriction, further studies will be necessary to understand the causes of total or partial source limitations during seed filling among H11 plants.

Arabidopsis, rice, and maize plants overexpressing *HaHB11* set more seeds than their controls (Cabello *et al*., 2016; Raineri *et al*., 2019; Raineri *et al*., 2022; Raineri *et al*., 2023). However, each of these transgenic species exhibited particular traits. Rice H11 plants showed a more erect architecture and increased partitioning to grains under normal growth conditions (Raineri *et al*., 2023) and maize H11 outyielded controls across different genetic backgrounds (B72 lines or B73 x Mo17 hybrids), grown under normal and stress conditions, such as flooding and defoliation (Raineri *et al*., 2022). To date, we have not identified a specific molecular pathway regulated by HaHB11 explaining such robust traits in different plant species, except of that high expression levels of the transgene are needed (Cabello *et al*., 2016; Raineri *et al*., 2022). Fig. 7 summarizes the differential traits between H11 and control plants we detected along with soybean life cycle.

**Figure 7.**
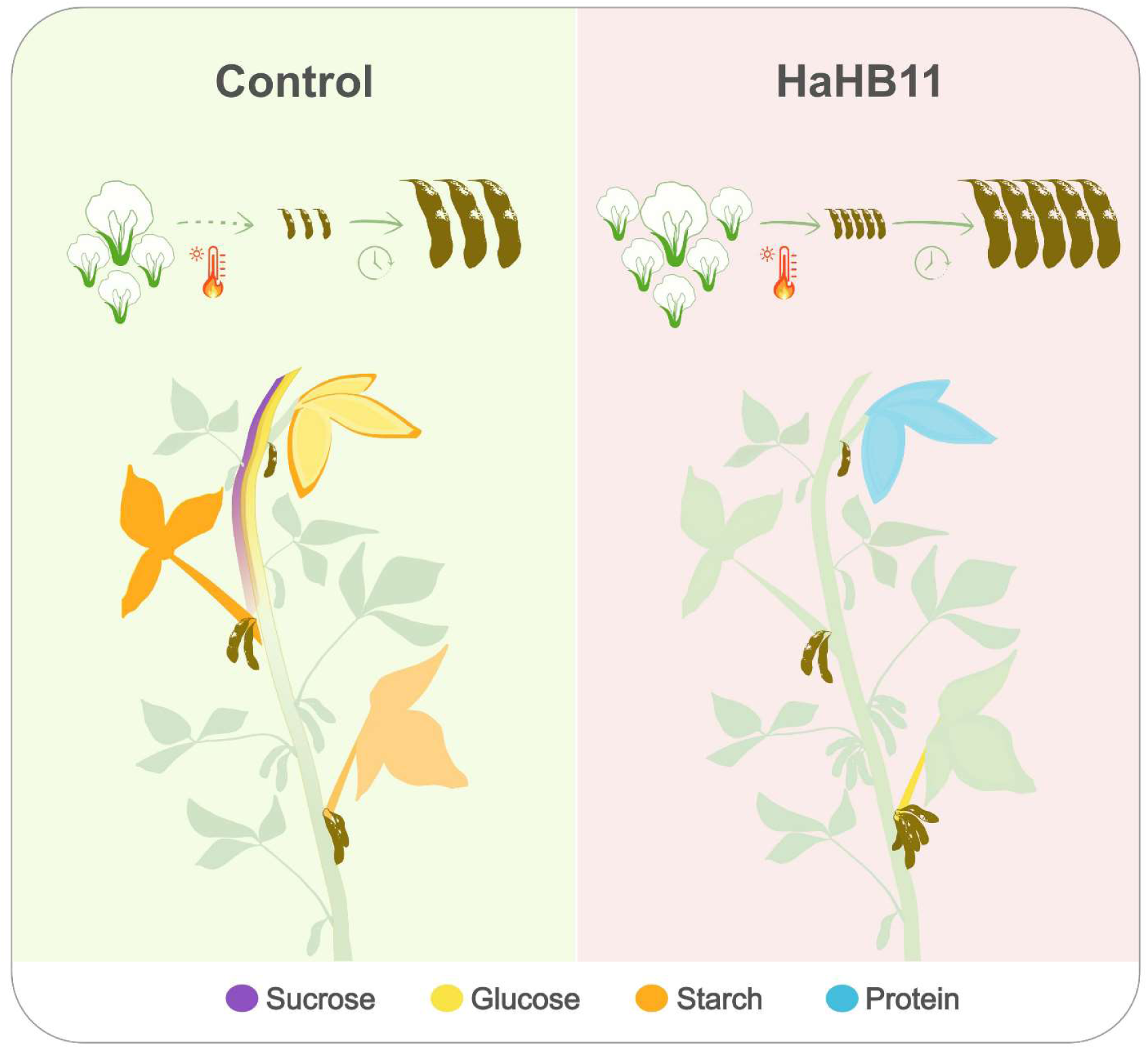
Schematic representation of the differential traits between H11 and control plants. Up: H11 flowers generating more pods than controls under heat stress, process taking more time but very synchronized. Middle: H11 plants have less carbohydrates than controls and produce a higher number of pods in the basal to mid-segment of the plant. The contents of sucrose (red), glucose (yellow), starch (orange), and protein (light blue) are represented in nodes 5, 8, and 15.

In this study, we considered vegetation indices (VIs) and proximal reflectance measurements to differentiate transgenic soybean plants from their wild types. Breunig *et al*. (2011) highlighted the importance of considering crop phenology for soybean varieties classification. Genotypic proximity and management with the same sowing date minimized morphological and phenological differences, making cultivar differentiation more difficult. Crusiol *et al*. (2021) succeeded in classifying several soybean cultivars via linear discriminant analysis, but they had higher errors in discriminating cultivars of the same genetic background. VIs with weak to moderate correlation were identified, ensuring no redundancy between them to differentiate H11 from control plants. Other research linking spectral data to physiological characteristics of soybeans, such as yield (Galvão *et al*., 2009) or pod formation percentage (Djanaguiraman *et al*., 2019), illustrates the growing potential of spectral technologies in plant breeding research.

## Concluding remarks

The overexpression of the sunflower transcription factor HaHB11 triggers an increased grain setting in soybeans. This trait can be explained by an augmented number of pollinated flowers, even under heat stress. H11 pods also developed more synchronously, allowing them to set more pods. Given the lower contents of starch and sucrose in leaves, petioles, and stems, it can be concluded that high demand for assimilates is exerted on source tissues, which may cause a total trade-off between grain yield components in certain environments. This response demands further investigations. Altogether, the results allow us to conclude that HaHB11 is a biotechnological tool to improve soybean yield, particularly under heat stress.

## Acknowledgements

We thank Ayelén Rapoport for her professional assistance in the procurement of CONABIA and INASE (Ministry of Agriculture) permits. We are very grateful for the technical assistance provided by Mr. Manuel Franco.

## Authors’ contribution

Conceived and designed the experiments: JR, MEO, and RLC. Performed the experiments: JR, and EMB. Analyzed the data: JR, EMB, MP, MEO, and RLC. MC transformed soybean plants. Wrote the paper: JR, MP, MEO, and RLC.

## Supplementary Material

**Supplementary Figure S1:** Soybean plants expressing the sunflower *HaHB11* gene exhibit differential traits compared to their controls in four field assays

**Supplementary Figure S2:** Grain yield is correlated with grain number

**Supplementary Figure S3:** Flowers’ developmental stages differ between transgenic and control plants in the reproductive stage

**Supplementary Figure S4:** Pod number differs between H11 transgenic and control plants but not seeds per pod nor seed composition

**Supplementary Figure S5:** Transgenic plants accumulate more biomass than controls but do not differ in harvest index

**Supplementary Figure S6:** H11 and H4 fill pods more synchronously than controls

**Supplementary Table S1:** H11 plants outyield controls in the seasons having high temperatures between R3-R5.

